# Can a bulky glycocalyx promote catch bonding in early integrin adhesion? Perhaps a bit

**DOI:** 10.1101/2023.03.16.532909

**Authors:** Aaron Blanchard

## Abstract

Many types of cancer overexpress bulky glycoproteins to form a thick glycocalyx layer. The glycocalyx physically separates the cell from its surroundings, but recent work has shown that the glycocalyx can paradoxically increase adhesion to soft tissues and therefore promote the metastasis of cancer cells. This surprising phenomenon occurs because the glycocalyx forces adhesion molecules (called integrins) on the cell’s surface into clusters. These integrin clusters have cooperative effects that allow them to form stronger adhesions to surrounding tissues than would be possible with equivalent numbers of un-clustered integrins. These cooperative mechanisms have been intensely scrutinized in recent years; a more nuanced understanding of the biophysical underpinnings of glycocalyx-mediated adhesion could uncover therapeutic targets, deepen our general understanding of cancer metastasis, and elucidate general biophysical processes that extend far beyond the realm of cancer research.

This work examines the hypothesis that the glycocalyx has the additional effect of increasing mechanical tension experienced by clustered integrins. Integrins function as mechanosensors that undergo catch bonding – meaning the application of moderate tension increases integrin bond lifetime relative to the lifetime of integrins experiencing low tension. In this work, a three-state chemomechanical catch bond model of integrin tension is used to investigate catch bonding in the presence of a bulky glycocalyx.

This modeling suggests that a bulky glycocalyx can lightly trigger catch bonding, increasing the bond lifetime of integrins at adhesion edges by up to 100%. The total number of integrin-ligand bonds within an adhesion is predicted to increase by up to ~60% for certain adhesion geometries. Catch bonding is predicted to decrease the activation energy of adhesion formation by ~1-4 k_B_T, which translates to a ~3-50× increase in the kinetic rate of adhesion nucleation. This work reveals that integrin mechanic and clustering likely both contribute to glycocalyx-mediated metastasis.

**Graphical Abstract:** 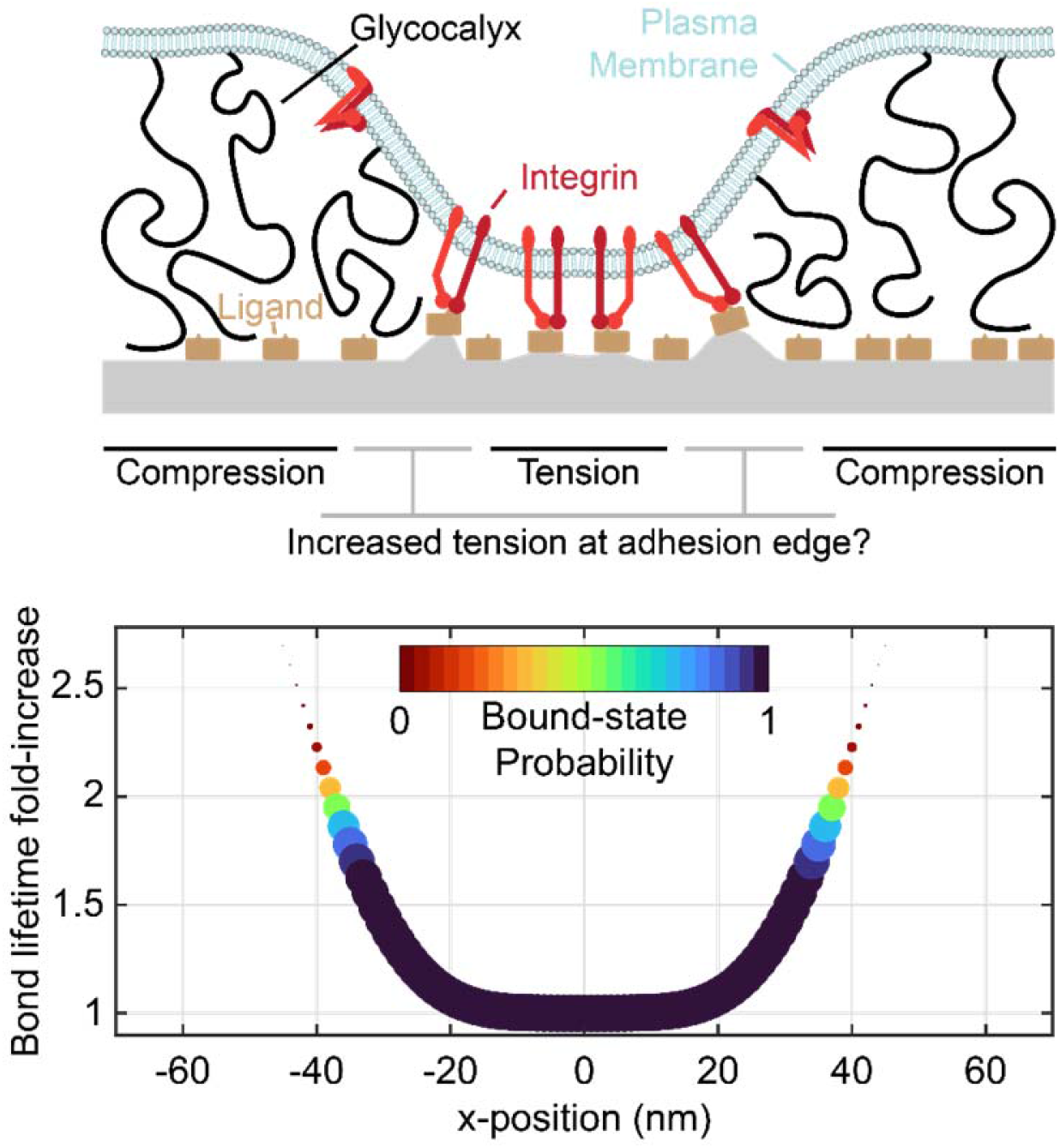

## Introduction

Many metastatic carcinomas – such as recurrent glioblastoma multiform, which is invariably lethal^1,2^ – overexpress bulky glycoproteins to form a thick (~10s-100s of nanometers [nm]) glycocalyx layer^3–6^. This glycocalyx imposes a physical separation between the cancer cell and its surroundings that is much wider than the typical length of adhesion proteins such as integrins (~20 nm). Recent work by Paszek, Weaver, and others has shown (perhaps counterintuitively) that this glycocalyx can *increase* adhesion to soft tissues and therefore *promote* the invasion of metastatic cancer cells by forcing integrin receptors into kinetic traps. Within these traps, the integrins assemble into high-strength focal adhesions (FAs) with the extracellular matrix (ECM)^6–8^ (Figure 1). FA formation and maturation is critical to invasion, survival, and growth of cancer cells; in addition to mechanically coupling the cytoskeleton and cell membrane to the ECM, FAs recruit signalling molecules that produce biochemical outputs^9,10^ including growth factor signalling upregulation^11,12^, which is also crucial for cancer proliferation^13–16^. In GBM (and potentially in similar types of soft-tissue cancer), integrin activation upregulates glycocalyx expression, leading to a feedback loop that further promotes cancer cell adhesion, survival, and growth^3^. Beyond GBM, glycocalyx upregulation is a common feature in metastatic and circulating cancer cells^4,6,17–20^, and therapies that de-bulk the glycocalyx have proven promising *in vivo*^21–23^.

**Figure 1:**
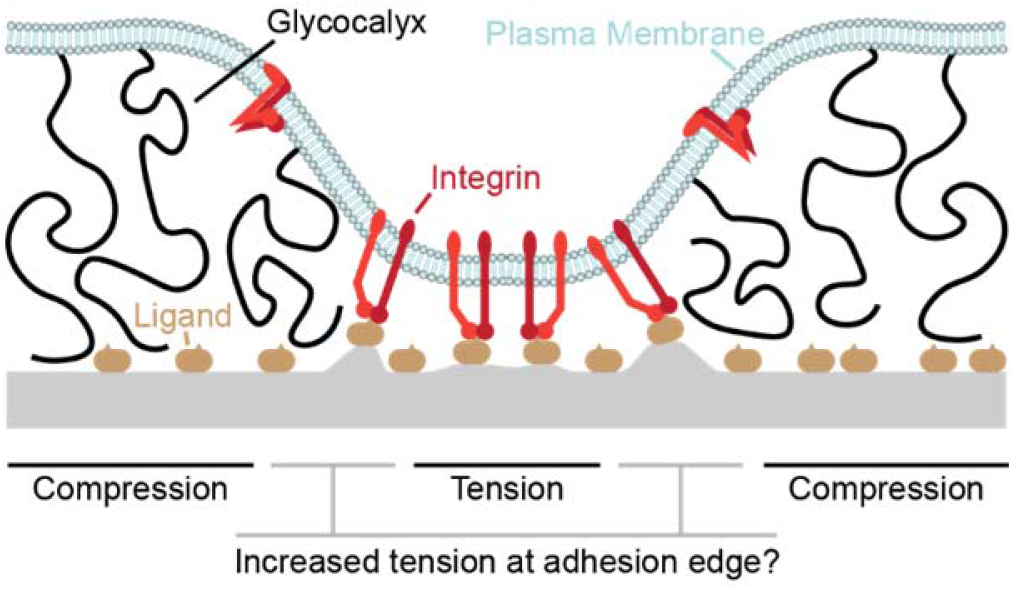
Schematic depiction of integrin adhesion and the glycocalyx

While recent work has illuminated the role that the glycocalyx plays in integrin *clustering*, much remains unknown about integrin *mechanics* in this context. Mechanical tension is critical to integrin function; integrins display catch-bond behavior, meaning that their adhesiveness increases under piconewton (pN)-scale tension^24^. Tension also promotes integrin structure-switching from an inactive conformation into an active conformation that promotes heightened intracellular signaling^25–27^. Furthermore, recent studies have shown that integrins generally behave as orientation-dependent mechanosensors^25,28–30^, meaning that the geometry of glycocalyx-mediated FAs may play a unique role in mediating integrin activation that is separate from clustering. The role that the glycocalyx plays in modulating the magnitude and orientation of tension experienced by integrins, and the effect that such modulation will have on integrin catch bonding, has not been directly investigated.

It recent years, it has been suggested on numerous occasions that a bulky glycocalyx can impose tensile forces on integrins^31,32^, which could potentially drive catch bonding and integrin structure switching. This hypothesis of glycocalyx mediated integrin tension – that is, the hypothesis that resistance from a bulky glycocalyx on the plasma membrane can increase the tension experienced by integrin receptors binding to extracellular ligands – has not (to this author’s knowledge) been investigated experimentally or computationally. However, this prediction can be derived from a simple force-balance analysis of integrin adhesion (**Figure 1**); if compression of the glycocalyx is counter-balanced by tension on integrin-ligand bonds then the edges of adhesions, where the plasma membrane slopes away from the underlying surface, should exhibit increased integrin tension due to increased distance between the plasma membrane and the surface.

Several computational models of integrin adhesion have incorporated mechanical considerations of the glycocalyx^7,33–36^. Other computational models have incorporated consideration of integrin catch bonding^37–41^. In this work, the mechanics of both glycocalyx compression and integrin catch bonding are considered together to understand how these phenomena interact to influence early integrin adhesion. Chemomechanical modeling^42,43^ is used to address these questions.

The results of this analysis suggest that a bulky glycocalyx can trigger a small amount of catch bonding, increasing the bond lifetime of integrins at adhesion edges by up to 100%. This increase in integrin lifetime can increase the total number of integrin-ligand bonds within an adhesion by up to ~60% (depending on adhesion geometry). Finally, the ability of integrins to form catch bonds (instead of slip bonds) is predicted to decrease the activation energy of adhesion formation by ~1-4 k_B_T, which translated to a ~3- to ~50-fold increase in the kinetic rate of adhesion nucleation. While these values may not be quantitatively accurate due to assumptions that were made in the construction of the model used in this work, the general conclusion of this study is that glycocalyx repulsion and integrin catch bonding may interact to impose a small – but non-negligible – increase in adhesiveness and the rate of integrin adhesion formation. Following initial adhesion, glycocalyx-mediated integrin clustering and intracellular signaling, as described previously^3,6,7,16^, can trigger positive feedback loops to promote endurant adhesion and tissue invasion. This work contributes to our understanding of the mechanical role of the glycocalyx in early cancer adhesion.

A plasma membrane (light blue) is shown, deformed by a combination of integrin-ligand adhesion and glycocalyx-mediated repulsion. While the glycocalyx regions experience compression and the central adhesive region experiences tension, this work is concerned with the enhanced tension that should be experienced at the edges of the adhesive region.

## Methods

In the model used for this work, interactions are simulated between integrins on a plasma membrane and ligands on an underlying planar surface. The ligands are fixed in position, but the integrins can diffuse freely across the plasma membrane. The values of all parameters used in this study are listed in Table 1.

**Table 1:** Model Parameters (Used Unless Otherwise Stated)

| Name | Symbol | Value | Reference |
| --- | --- | --- | --- |
| Ligand surface density | $\rho_{ligand}$ | 2,500 $\mu\text{m}^{-2}$ | 7 |
| Active integrin surface density | $\rho_{integrin}$ | 10 $\mu\text{m}^{-2}$ | 7 |
| Integrin-ligand spring constant | $\kappa_{bond}$ | 2 pN/nm | 7 |
| Reactive compliance | $\gamma$ | 0.4 nm | 7 |
| Glycocalyx spring constant | $\kappa_{glyc}$ | 0.02 pN/nm | 7 |
| Plasma membrane grid spacing | $\delta_{PM}$ | 1 nm | Selected |
| Ligand grid spacing | $\delta_{Lig}$ | 1 nm | Selected |
| Prefactor (unbound bound<sub>1</sub>) | $k_{on,0}$ | 10<sup>5</sup> s<sup>-1</sup> | 7 |
| Prefactor (bound<sub>1</sub> to bound<sub>2</sub>) | $k_{1,0}$ | 1.185 s<sup>-1</sup> | Optimized<sup>a</sup> |
| Transition state distance (bound<sub>1</sub> to bound<sub>2</sub>) | $\Delta x_1$ | 0.342 nm | Optimized<sup>a</sup> |
| Prefactor (bound<sub>2</sub> to bound<sub>1</sub>) | $k_{-1,0}$ | 1.971 s<sup>-1</sup> | Optimized<sup>a</sup> |
| Transition state distance (bound<sub>2</sub> to bound<sub>1</sub>) | $\Delta x_{-1}$ | -0.626 nm | Optimized<sup>a</sup> |
| Prefactor (bound<sub>1</sub> to unbound) | $k_{off,1,0}$ | 2.004 s<sup>-1</sup> | Optimized<sup>a</sup> |
| Transition state distance (bound<sub>1</sub> to unbound) | $\Delta x_{off,1}$ | 0.427 nm | Optimized<sup>a</sup> |
| Prefactor (bound<sub>2</sub> to unbound) | $k_{off,2,0}$ | 0.021 s<sup>-1</sup> | Optimized<sup>a</sup> |
| Transition state distance (bound<sub>2</sub> to unbound) | $\Delta x_{off,2}$ | 0.204 nm | Optimized<sup>a</sup> |
| Slip bond prefactor | $k_{slip,0}$ | 1.075 s<sup>-1</sup> | Calculated |
| Slip bond transition state distance | $\Delta x_{slip}$ | 0.5 nm | Selected |
| Plasma membrane mean curvature stiffness | $\bar{\kappa}$ | 243 k<sub>B</sub>T/nm<sup>2</sup> | 45 |
| Plasma membrane Gaussian curvature stiffness | $\kappa_c$ | 243 k<sub>B</sub>T/nm<sup>2</sup> | 45 |
| Steady-state time | $t_{max}$ | 1,000 s | Selected |
| Simulation time step | $\Delta t$ | Variable | Variable |

<sup>a</sup> parameters provided by Asaro, Zhu, and Lin did not produce the expected catch bond behavior, so parameters were re-measured by tracing the $\tau$ vs. $F$ plot from that work and fitting parameters that best reproduced the traced curve.

### Kinetic and mechanical models of integrins

In line with prior work^7^, integrin ligand bonds in this work are modelled as linear springs with stiffness *κ_bond_* = 2 *pN/nm* (**Figure 2C)**. Accordingly, the relationships between bond force, *F*, mechanical strain energy (*G_bond_*), and bond extension (r) are described by the relationships:

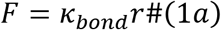

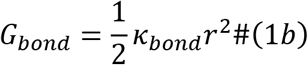

Integrin catch bonding is represented in this work using a three-state model, wherein the integrin-ligand pair undergoes force-dependent transitions between the unbound state and two distinct bound states (bound_1_ and bound_2_) as shown in **Figure 2A**^37,44^. Kinetic rate constants for transitions between states are modelled with Bell model^33^-type equations:

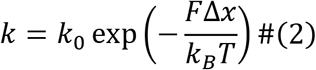

where *k*_0_ is the prefactor parameter, Δ*x* is the distance to the transition state parameter, and *k_B_T* = 4.114 *pN nm* is thermal energy at room temperature. Specifically, transitions from bound_1_ to bound_2_ and vice-versa (*k*_1_ and *k*_−1_, respectively) and from bound_1_ and bound_2_ to unbound (*k*_*off*,1_ and *k*_*off*,2_, respectively) are described by:

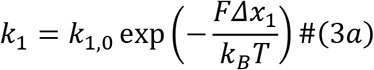

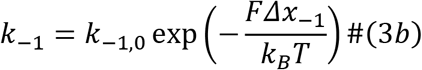

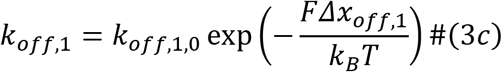

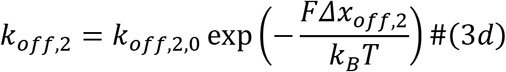

where the 8 Bell model parameters (**Figure 2D**) were optimized to recreate the average lifetime (*τ*) vs. *F* plot from Asaro, Lin, and Zhu^37^ as shown in (**Figure 2E**).

**Figure 2:**
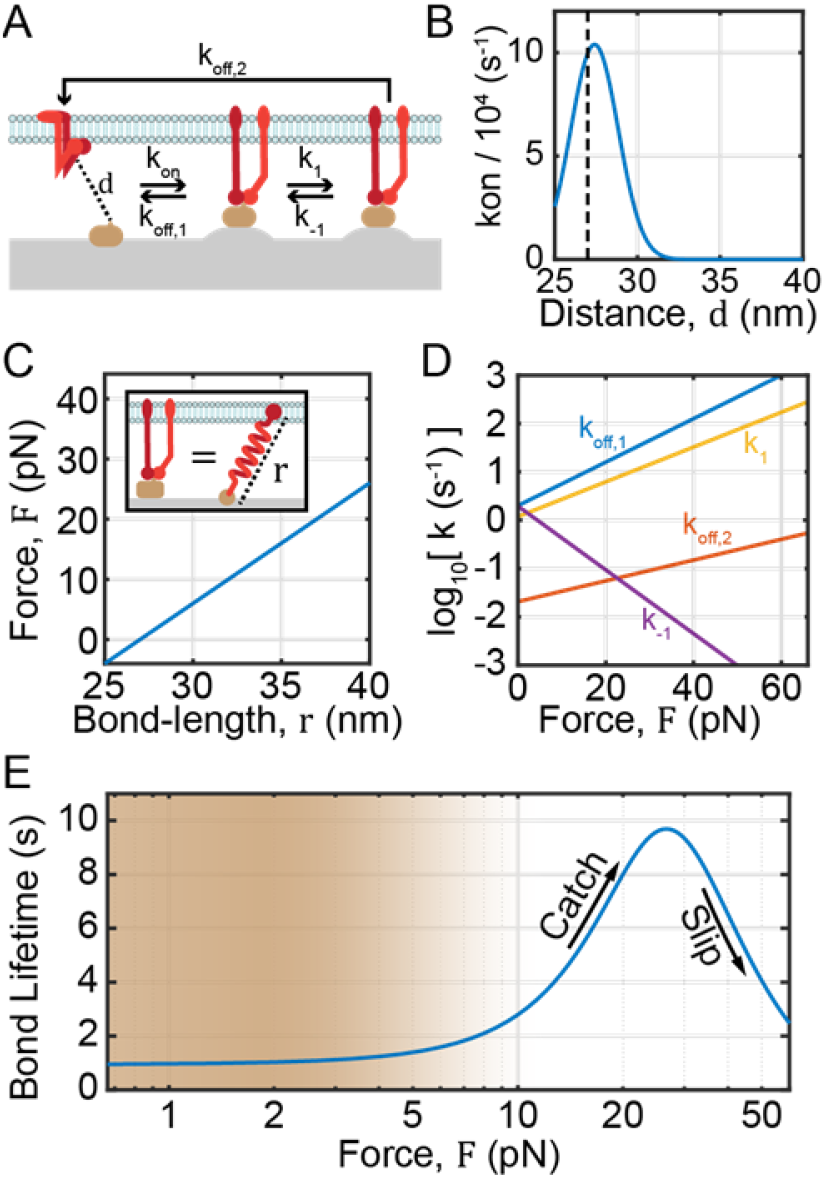
Mechanical and kinetic relationships used in this work. **A)** In this work, integrin catch bonding was modelled using a three-state model wherein integrins transition between unbound (left), loosely bound (or bound_1_, middle), and tightly bound (or bound_2_, right) states. The kinetic rate constants for transitions between these states are shown with flow arrows. **B)** The integrin-ligand association rate, *k_on_*, is modelled as a function of integrin-ligand pair distance, *d*, using a simple transition state energy model. Vertical dashed line shows the equilibrium integrin-ligand bond length. **C)** The relationship between integrin-ligand bond force, *F*, and extension, *r*, is modelled as a linear spring with stiffness *κ* = 2 *pN nm*^-2^ and a resting length of 27 *nm*. **D)** Additional kinetic rate constants for the optimized three-state model are shown as a function of *F* on a semi-logarithmic plot. **E)** Average bond lifetime on a semi-log scale as a function of force for the optimized three-state model. Shading denotes the general regime of forces studied in this work, and arrows show the catch and slip regimes of integrin forces.

Bond formation is modelled by transitions between the unbound state to the bound1 state, and occurs with a kinetic rate constant *k_on_* as a function of the integrin ligand distance, *d* (**Figure 2B**):

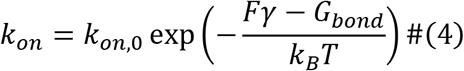

where *F* and *G_bond_* are calculated at r = *d* and *γ* = 0.4 *nm* is the reactive compliance parameter^6^.

To assess the importance of catch bonding in this work, results were also compared to results obtained using slip bond kinetics, represented with the equation:

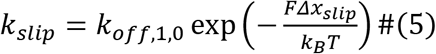

### Physical representation of the plasma membrane

The plasma membrane is approximated as a square grid, centered on the origin, with 1 nm lateral spacing and zero thickness. The plasma membrane is assumed to have a Gaussian profile (this shape was chosen because it resembles known profiles and can be described simply with 3 parameters). The z-height of the *i^th^* gridpoint, *z_PM,i_*, can thus be described by:

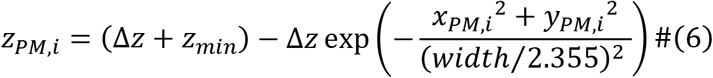

where *x_PM,i_* and *y_PM,i_* are the x- and y-positions of the i^th^ gridpoint. *z_min_* is the minimum height of the plasma membrane, Δz is the difference in the minimum and maximum height of the plasma membrane, and *width* is full-width half at half maximum of the Gaussian (e.g. the width of the plasma membrane protrusion at *z* = *z_min_* + Δ*z*/2) (**Figure 3C**). The shape of the adhesion is treated as static, maintained by a spatiotemporal ensemble of integrin-ligand bonds. The width of the full grid was set to 600 nm, which is substantially larger than the width values simulated in this work.

**Figure 3:**
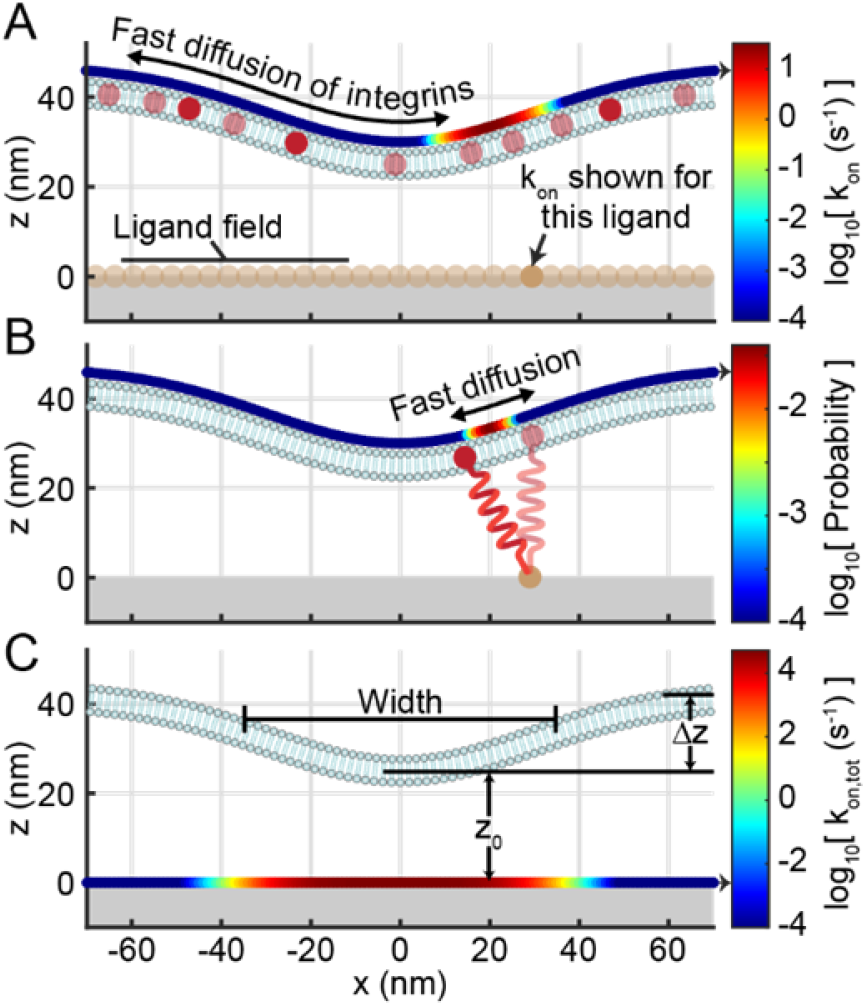
Representation of physical model studied in this work. **A)** A membrane adhesion is shown, with integrins (red dots) shown freely diffusing across it, along with a nearly-continuous field of ligands. Above the plasma membrane, a coloured curve shows *k_on_* between the ligand (denoted with an arrow) and each position on the plasma membrane. **B)** A single integrin-ligand bond, with free diffusion of the integrin along the plasma membrane, is shown. A curve shows the probability density function of the integrin’s position. **C)** The total association rate, *k_on,tot_*, is shown for all positions on the ligand field with a coloured curve on the underlying surface. Physical parameters Width, Δ*z*, and *z*_0_ are also depicted. In this figure, Width=165 nm, *z*_0_ = 25 *nm*, and Δ*z* = 18 *nm*.

The field of ligands on the planar surface was approximated as a one-dimensional grid with 1 nm spacing between gridpoints, gridpoint y-values (*y_S_*) of 0, z-values (*z_S_*) of 0, and x-values (*x_S_*) ranging from −600 to 600. The 1-dimensional grid is less computationally expensive than simulating a full 2-dimensional grid of surface points. Due to the radial symmetry of the adhesion used in the work, the result that would have been with a 2-dimensional surface grid can be recovered by integrating the 1-dimensional result radially about the origin.

### Steady-state calculation

To calculate the steady-state number of integrin-ligand bonds at a given surface position, total rate constants were first calculated by integration. The rate constant for total association at the j^th^ substrate gridpoint was calculated by numerically integrating *k_on_* between the substrate gridpoint and all plasma membrane gridpoints within a lateral cutoff of 50 nm:

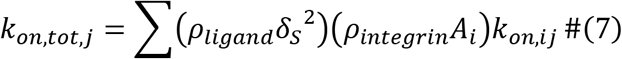

where *k_on,ij_* is *k_on_* calculated with *d* as the distance between the j^th^ substrate gridpoint and the i^th^ plasma membrane gridpoint, *ρ_ligand_* is the surface density of ligands on the planar surface, *ρ_integrin_* is the surface density of active integrins on the plasma membrane (**Figure 3A, C**). *A_i_* is the area of the i^th^ gridpoint, which is can be approximated by:

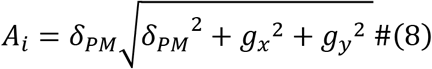

where *g_x_* and *g_y_* are the x- and y-components of the gradient of the plasma membrane surface.

Each of the other rate constants for the j^th^ gridpoint was also calculated:

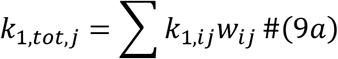

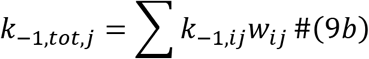

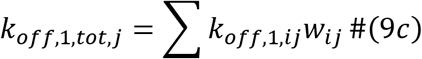

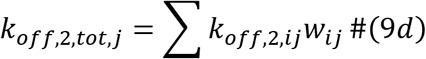

where *w_ij_* is the fraction of time that an integrin paired to the ligand at the j^th^ substrate gridpoint spends at the i^th^ plasma membrane gridpoint (**Figure 3B**). *w_ij_* was calculated using a partition function:

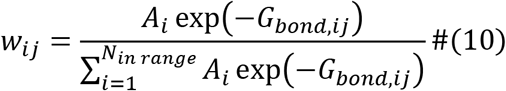

Once all total rate parameters are calculated, an Eulerian Markov matrix, ***M***, was constructed:

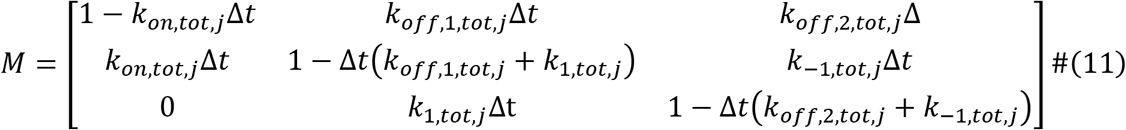

where Δ*t*, the simulation timestep, is calculated for numerical stability. Finally, steady-state calculation of the fraction of time the ligand at the j^th^ gridpoint spends in the unbound state (*f_unbound_*), in the bound_1_ state (*f*_*bound*,1_), and in the bound_2_ state (*f*_*bound*,2_) is performed via matrix multiplication:

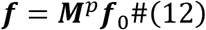

where *p* = *t_max_*/Δ*t*, *t_max_* is the steady-state time (1,000 *s*), ***f*** = [*f_unbound_ f*_*bound*,1_ *f*_*bound*,2_]^T^ (^*T*^ denotes transposition), and ***f***_0_ = [1 0 0]^*T*^ denotes zero initial adhesion.

### Energetics

Adhesion energy was calculated as the number of integrin-ligand bonds multiplied by the free energy per bond (5 k_B_T). The summed mechanical strain energy of the integrin ligand bonds was not factored into this calculation because it is intrinsically factored into the steady-state calculation of the number of bonds.

Plasma membrane bending energy was calculated by summing the Helfrich^45^ energy (assuming zero spontaneous curvature) across all gridpoints:

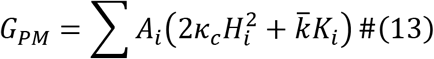

where *H_i_* and *K_i_* are the Gaussian and mean curvatures (calculated using the “surfature” command, shard on the MathWorks file exchange by D. Claxton), respectively, of the i^th^ gridpoint and *κ_c_* and 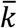 are bending stiffness parameters. Finally, the free energy of glycocalyx compression is calculated by integrating the mechanical strain energy of the glycocalyx across all gridpoints:

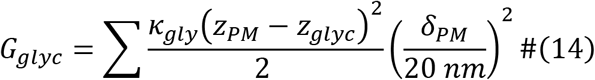

where *κ_glyc_* is the spring constant of the glycocalyx, *z_glyc_* = 43 *nm* is the glycocalyx equilibrium length, and 20 *nm* is the gridpoint spacing used by Paszek *et al*.^7^.

## Results and Discussion

### Initial representative calculation

Initial steady-state calculations of integrin adhesion were performed (using parameters from previous work^7^) with *Width* = 165 *nm*, *z*_0_ = 27 *nm*, and Δ*z* = 16 *nm*. For comparison as a glycocalyx-free control, steady-state adhesion to a flat plasma membrane was also simulated (*z*_0_ = 27 *nm* and Δ*z* = 0 *nm*) (**Figure 4A,B**). The flat plasma membrane resulted in a high degree of adhesion across the surface, with *f_unbound_* ≈ 0, *f*_*bound*,1_ ≈ 0.55, and *f*_*bound*,2_ ≈ 0.45 (**Figure 4C**). In contrast, binding to the curved plasma membrane was highly position-dependent; *f_unbound_* increased from 1 far away from the center of the adhesion to ~0 close to the center of the adhesion (**Figure 4D**). Notably, the bound_2_ state (the tightly bound state) was higher at the edge of the plasma membrane contact zone (peaking at ~0.85) than at the center of the contact zone.

**Figure 4:**
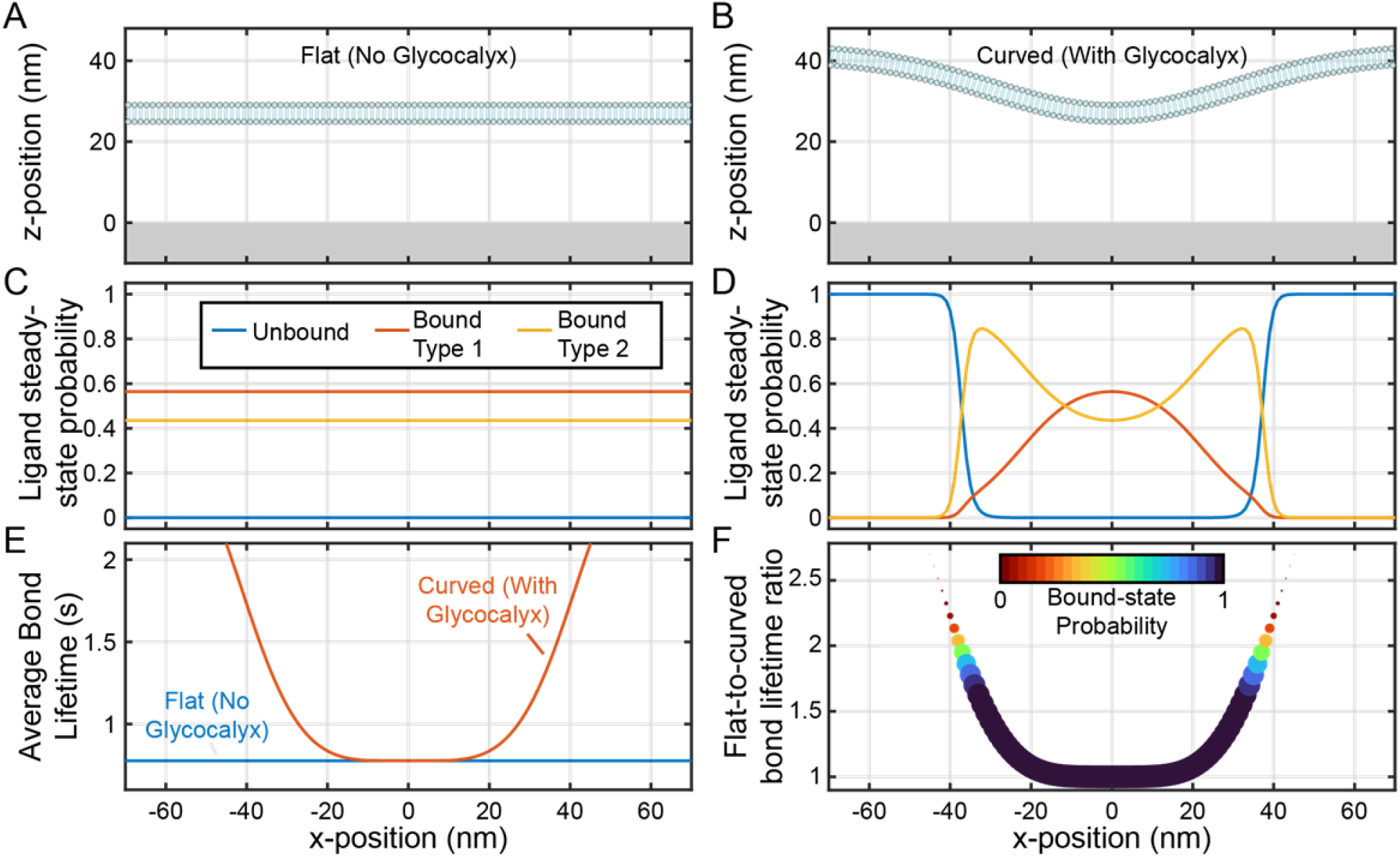
A bulky glycocalyx promotes catch bonding and slightly increases bond lifetime at adhesion edges. **A)** Physical representation of a flat plasma membrane and **B)** a curved plasma membrane. Both have the same minimum position (*z*_0_ = 27 *nm*), but the curved plasma membrane curves up to a height of 43 nm (Δ*z* = 16 *nm*). **C)** Probabilities that ligands beneath the plasma membrane are in the unbound, loosely bound, and tightly bound states for the flat plasma membrane, shown as a function of ligand x-position. **D)** Same as **C**, but for the curved plasma membrane. **E)** The average bound lifetime as a function of ligand x-position for both the curved and flat plasma membranes. **F)** Bound lifetime for the curved membrane divided by the bound lifetime for the flat membrane, with the probability of existing in a bound state depicted with both color and marker size, as a function of ligand x-position.

The substantial increase in the tightly bound state observed for the curved plasma membrane provides support for the hypothesis that the glycocalyx promotes catch bonding. To better assess the catch bond-mediated lifetime increase, the average bond lifetime at each surface gridpoint was calculated be re-arranging the equilibrium constant equation:

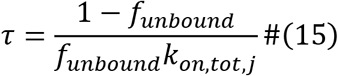

While *τ* was low for the flat plasma membrane (~0.93 s), *τ* increased up as high as 2 seconds at the edge of the contact zone. The ratio of *τ* with a flat plasma membrane to *τ* with a curved plasma membrane, *θ*, which is equivalent to the increase in bond lifetime that is driven by the presence of a bulky glycocalyx, increases as high as 2.5 (**Figure 4F**). However, these high *θ* values were concentrated at positions with high *f_unbound_* values. Some locations exhibited both nonzero binding and a high *θ;* for example, at *x_S_* = ±35 *nm*, 1 – *f_unbound_* ≈ 0.1 and the *θ* ≈ 2.

The catch binding-mediated percent-increase in binding across the entire adhesion, *ϵ*, can be calculated through integration by weighting according to the bound fraction at each surface gridpoint:

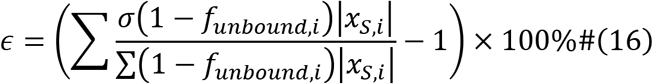

For this initial calculation, *ϵ* = 27.7% which is a moderate increase in adhesion due to catch-bonding and the presence of a bulky glycocalyx.

### Binding enhancement with different adhesion geometries

Calculations of *ϵ* were next performed across a range of adhesion geometries. Specifically, *z*_0_ was varied from 20 nm (representing very close contact) to 40 nm (representing a complete lack of binding). Width was varied from 5 nm to 240 nm. For all conditions, the maximum plasma membrane height, *z*_0_ + Δ*z* was held fixed at *z_glyc_* = 43 *nm*. In addition, the total number of integrin-ligand bonds, *n_bond_* was calculated by numerical integration:

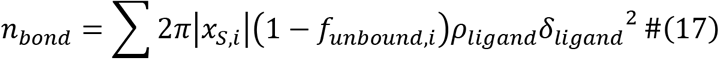

For this initial set of calculations, the maximum ligand density simulated by Paszek *et al.^7^* was used (*ρ_ligand_* = 2,500 *μm*^-2^). Across all *z*_0_ values, *n_bond_* increased superlinearly with adhesion width, while *ϵ* increased slightly with increasing width and levelled off around *width* ≈30 nm (**Figure 5A**). In contrast, *n_bond_* decayed to zero and *ϵ* increased superlinearly with increasing *z_0_* across all *width* values (**Figure 5B**). To summarize, *n_bond_* depends strongly on both *z*_0_ and *width* (**Figure 5C**), but *ϵ* depends primarily on *z*_0_ (**Figure 5D**).

**Figure 5:**
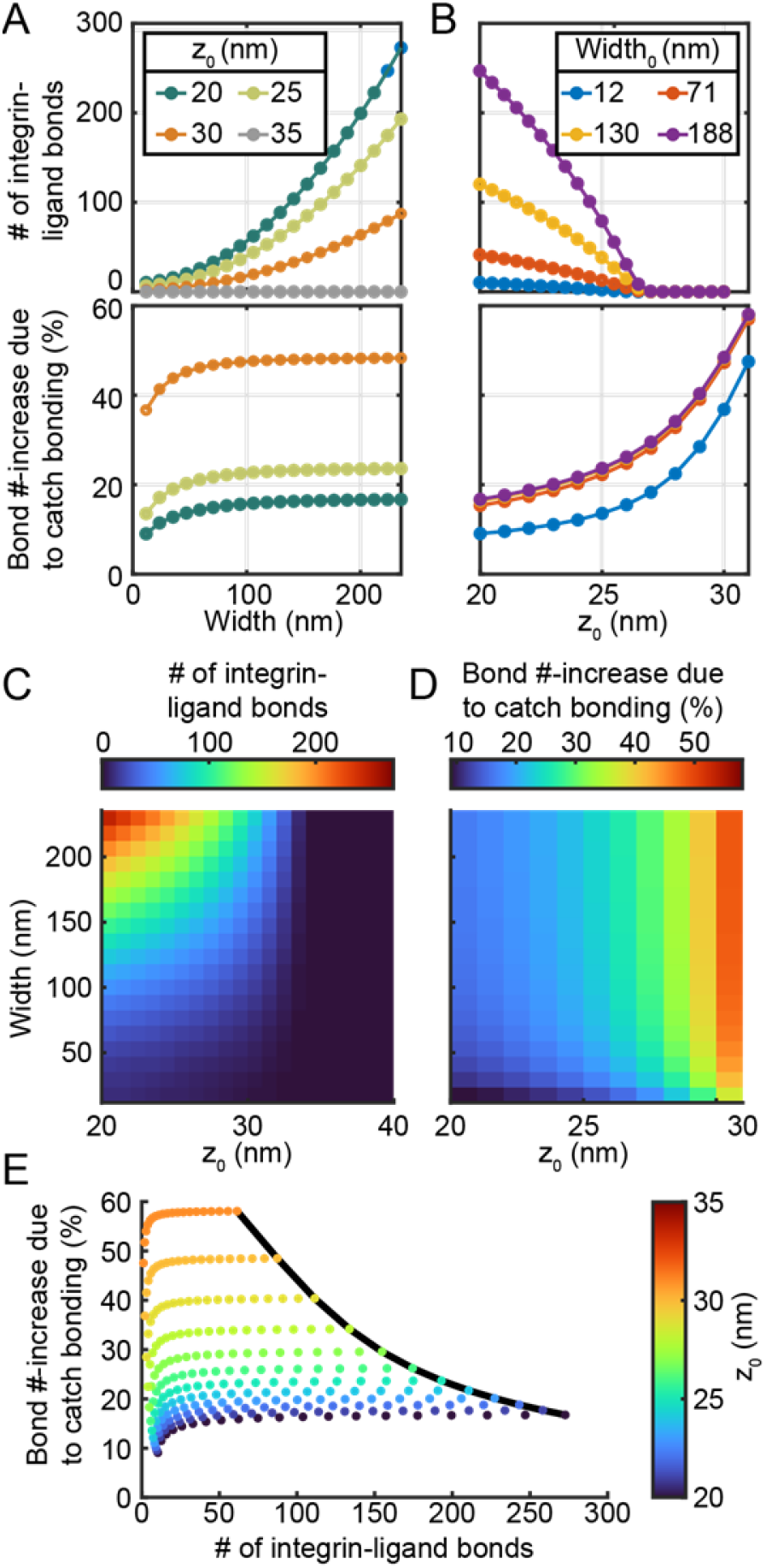
Glycocalyx- and catch bond-mediated adhesion enhancement as a function of adhesion geometry. **A)** The total, steady-state number of integrin bonds (top) and percent-increase in the total number of bonds when catch bonding is enabled (compared to slip-bonding) (bottom) as a function of adhesion width at four different *z*_0_ values. Percent increase is not shown for *z*_0_ = 35 *nm* due to negligible binding at this condition. **B)** Same as **A**, but as a function of *z*_0_ at 4 different adhesion widths. **C)** Surface plots showing the total bond number and **D)** bond number percent-increase as functions of *z*_0_ and Width. **E)** Scatter plot of bond number percent-increase vs. total bond number for all conditions tested in **C** and **D**. Scatter point color indicates *z*_0_, while width increases from left to right. A black curve shows the boundary; generally, the highest percent-increases occur at low total bond numbers, suggesting that the total effect of integrin catch bonding is limited and constrained across the entire parameter space.

As before, the largest *ϵ* is observed at conditions that have the least amount of binding (e.g. at higher *z*_0_). This finding suggests an overall limit to the enhancement effect. Indeed, at conditions with 1 – *f_unbound_* > 0.001 (that is, when *z*_0_ < 30 *nm*), the frontier of the *ϵ* vs. *n_bond_* scatterplot (black line, which occurs at *width* = 236 *nm*, **Figure 5E**) fits moderately well (R^2^=0.925) to the equation *ϵ* = 41.32/*n_bond_*. Re-arranging this equation reveals that, at *width* = 236 *nm*, *n_catch_* =41.32 integrin-ligand bonds are gained on average due to catch bonding, regardless of *z*_0_ in this range. Within this same range of *z*_0_ values, the number of bonds gained due to catch bonding, *n_catch_*, increased linearly with the adhesion surface area; when *z*_0_ < 30 *nm*, *n_catch_* fit well (R^2^=0.99) to the relationship *n_catch_* = (8.13 × 10^-4^)(width/nm)^2^.

The process of initial adhesion formation is characterized primarily by decreases of *z*_0_ (e.g., the tip of adhesion coming into close contact with the substrate). The process of adhesion growth is characterized by increasing *Width*. The impact of catch binding on both processes can be analysed separately. The results of the previous paragraph show that during the adhesion growth process, catch bonding continues to play an important role in increasing binding at the adhesion edge. However, as the adhesion grows, the total fraction of integrin-ligand bonds that exhibit catch binding decreases (because bonds that are not close to the adhesion edge experience lower intrinsic forces).

### Effect of catch bonding on adhesion formation

To assess the dynamics of initial adhesion formation, the system energetics were considered by summing the energies of glycocalyx compression (**Figure 6A, B**), membrane bending (**Figure 6C**), and adhesion (**Figure 6D**) (see **Methods** subsection “Energetics”):

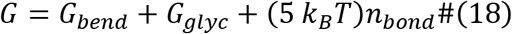

A phase plane analysis was then applied to the *G* vs. *width* vs. *z*_0_ surface (**Figure 7A**), which contained one saddle point. Gradient descent was used to determine that the energy surface exhibits two regions: One region wherein adhesion growth is favored, and one region wherein adhesion shrinkage is favored. At moderate and high *width* (i.e. >~70 nm), the dividing line (i.e. the separatrix, blue line in **Figure 7A**) between these two regions occurs at *z*_0_ = 30 to 35 *nm*. Very narrow-width adhesions are energetically disfavoured due to high plasma membrane bending energy, and so the separatrix shifts towards *z*_0_ ≈ 25 *nm* at low *width*.

**Figure 6:**
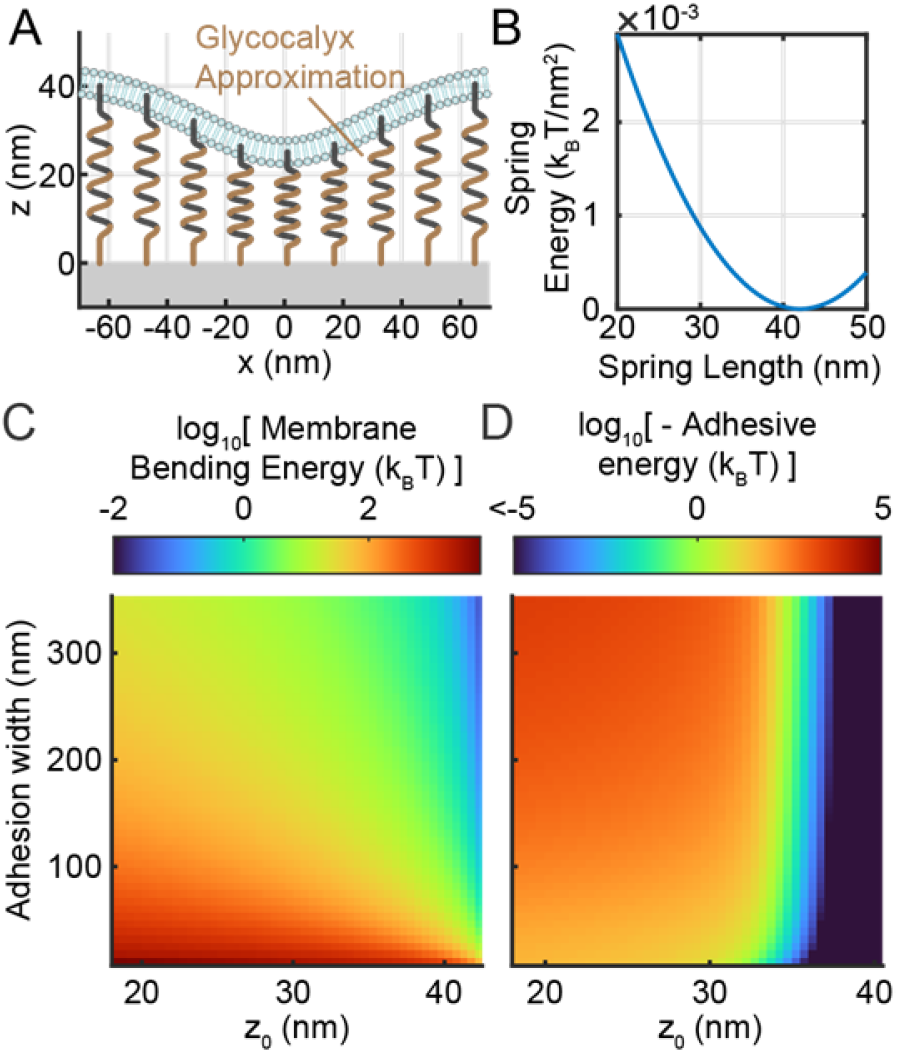
Energetics of glycocalyx compression, membrane bending, and adhesion. **A)** Representation of the glycocalyx as a layer of springs. **B)** Spring energy of the glycocalyx layer as a function of spring length. **C)** Total membrane bending energy and **D)** steady-state energy of integrin ligand adhesion as a function of adhesion width and Δ*z*.

**Figure 7:**
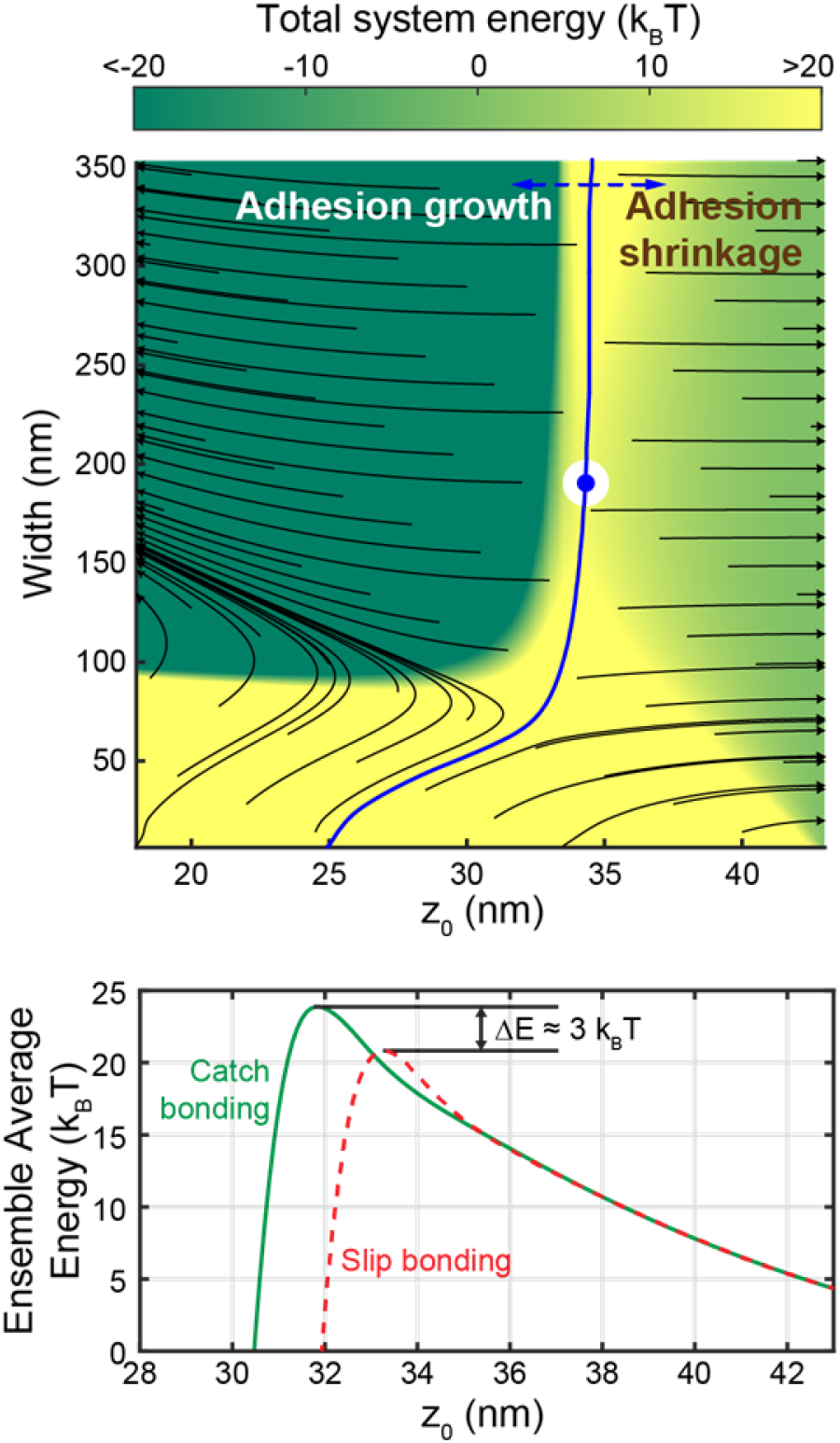
System energetics of adhesion

This energy surface was then used to calculate the activation energy of adhesion formation. To accomplish this, the ensemble average energy at each *z*_0_ value was calculated using the partition function:

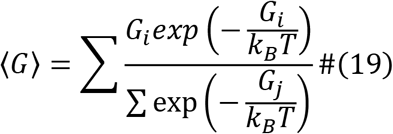

Calculating 〈*G*〉 at each *z*_0_ value yields a collapsed energy vs. *z*_0_ curve (**Figure 7B**) representing the process of initial adhesion formation. In a manner resembling the energy surface, the collapsed curve exhibits a single maximum that separates the adhesion shrinkage and adhesion growth regimes. In order to assess the role of catch bonding in promoting adhesion formation, this process was also repeated with integrins that exhibit slip-bonding (**Figure 7B**). A 1.5 nm shift in the location of the energy maximum, as well as a Δ*E* ≈3 k_B_T decrease in the height of the maximum, can be seen in the resulting energy vs. *z*_0_ curve. This change in the activation energy barrier for adhesion formation translates to an exp(3) ≈ 20-fold change in the kinetic rate of adhesion formation.

Surface plot (top) showing total system energy as a function of adhesion width and *z*_0_. Black flow lines are shown, as well as blue separatrix curve and a blue and white marker showing the surface’s energy plot. Adhesion growth is favored to the left of the separatrix, while adhesion shrinkage is favored to the right of the separatrix. Ensembled averaged energy vs. *z*_0_ plots (bottom) are shown for catch bonding and slip bonding. The activation energy difference between catch and slip bonding is shown as 3 multiples of thermal energy.

To conclude this study, this process was repeated to compute ΔE at lower ligand densities (*ρ_ligand_*, ranging from 250 to 2,500 *μm*^-2^) and different bond stiffnesses (*κ_bond_*) (**Figure 8**). Lower bond densities are expected to be more physiologically relevant than what was originally simulated, while lower bond stiffness can serve as an approximation for adhesion with compliant materials (such as soft tissue). This analysis suggests that Δ*E* is independent of *ρ_ligand_* (**Figure 8A**) but depends nearly linearly on the *κ_bond_* (**Figure 8B**). In other words, this analysis suggests that the role of integrin catch bonding in early adhesion is independent of ligand density, but dependent on bond stiffness. The Δ*E* values in this analysis ranged from 1.5 to 3.5 k_B_T, corresponding to a 4.5× to 33× increase in the rate of adhesion formation.

**Figure 8:**
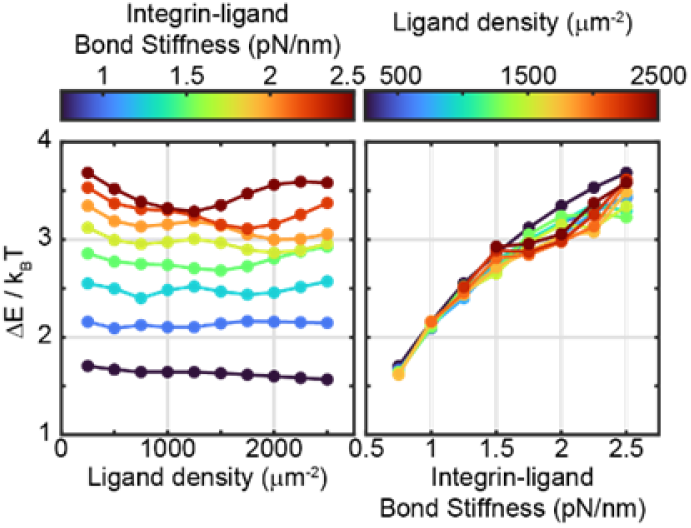
Catch bond-mediated change in activation energy (Δ*E*) as a function of ligand density (left) and integrin ligand bond stiffness, *κ* (right).

## Conclusion

This study suggests that catch bonding may perform a small role in promoting integrin adhesion due to intrinsic stress applied by the glycocalyx. Simulations in this work show that integrin catch bonding at the edges of curved adhesions can increase integrin-ligand bond lifetime by ~2-fold (**Figure 4**) and can increase the total number of integrin-ligand bonds by ~15-50% (**Figure 5**). Compared to hypothetical integrins with slip bond kinetics, integrins with catch-bond kinetics are predicted to form with a decrease in activation energy on the order of 1-4 k_B_T, corresponding to a 2.7-55× increase in the kinetic rate of adhesion formation. As with any computational study, numerous assumptions and simplifications were made that could alter these quantitative results. Nonetheless, the interaction between catch bond kinetics and a bulky glycocalyx in integrin function has not yet been investigated. This initial study suggests that catch bonding likely plays a small-but-notable role in early integrin adhesion in the presence of a bulky glycocalyx.

## Acknowledgments

This work was supported by the United States National Cancer Institute F99/K00 fellowship, grant number F99CA245789 / K00CA245789.

## References

1 Louis, D. N. et al. The 2016 World Health Organization Classification of Tumors of the Central Nervous System: a summary. Acta Neuropathologica 131, 803–820, doi: 10.1007/s00401-016-1545-1 (2016).

2 Gallego, O. Nonsurgical treatment of recurrent glioblastoma. Current oncology (Toronto, Ont.) 22, e273–281, doi: 10.3747/co.22.2436 (2015).

3 Barnes, J. M. et al. A tension-mediated glycocalyx–integrin feedback loop promotes mesenchymal-like glioblastoma. Nat. Cell Biol. 20, 1203–1214, doi: 10.1038/s41556-018-0183-3 (2018).

4 Pinho, S. S. & Reis, C. A. Glycosylation in cancer: mechanisms and clinical implications. Nature Reviews Cancer 15, 540, doi: 10.1038/nrc3982 (2015).

5 Wade, A. et al. Proteoglycans and their roles in brain cancer. The FEBS Journal 280, 2399–2417, doi: doi: 10.1111/febs.12109 (2013).

6 Paszek, M. J. et al. The cancer glycocalyx mechanically primes integrin-mediated growth and survival. Nature 511, 319, doi: 10.1038/nature13535 (2014).

7 Paszek, M. J., Boettiger, D., Weaver, V. M. & Hammer, D. A. Integrin Clustering Is Driven by Mechanical Resistance from the Glycocalyx and the Substrate. PLoS Comput. Biol. 5, e1000604, doi: 10.1371/journal.pcbi.1000604 (2009).

8 Woods, E. C., Yee, N. A., Shen, J. & Bertozzi, C. R. Glycocalyx Engineering with a Recycling Glycopolymer that Increases Cell Survival In Vivo. Angewandte Chemie (International ed. in English) 54, 15782–15788, doi: 10.1002/anie.201508783 (2015).

9 Hynes, R. O. Integrins: Bidirectional, Allosteric Signaling Machines. Cell 110, 673–687, doi: https://doi.org/10.1016/S0092-8674(02)00971-6 (2002).

10 Geiger, B. & Yamada, K. M. Molecular architecture and function of matrix adhesions. Cold Spring Harbor perspectives in biology 3, doi: 10.1101/cshperspect.a005033 (2011).

11 Cabodi, S. et al. Integrin regulation of epidermal growth factor (EGF) receptor and of EGF-dependent responses. Biochemical Society transactions 32, 438–442, doi: 10.1042/bst0320438 (2004).

12 Moro, L. et al. Integrin-induced epidermal growth factor (EGF) receptor activation requires c-Src and p130Cas and leads to phosphorylation of specific EGF receptor tyrosines. The Journal of biological chemistry 277, 9405–9414, doi: 10.1074/jbc.M109101200 (2002).

13 Shibue, T. & Weinberg, R. A. Integrin beta1-focal adhesion kinase signaling directs the proliferation of metastatic cancer cells disseminated in the lungs. Proceedings of the National Academy of Sciences of the United States of America 106, 10290–10295, doi: 10.1073/pnas.0904227106 (2009).

14 Desgrosellier, J. S. & Cheresh, D. A. Integrins in cancer: biological implications and therapeutic opportunities. Nature reviews. Cancer 10, 9–22, doi: 10.1038/nrc2748 (2010).

15 Freeman, S. A. et al. Integrins Form an Expanding Diffusional Barrier that Coordinates Phagocytosis. Cell 164, 128–140, doi: 10.1016/j.cell.2015.11.048 (2016).

16 Woods, E. C. et al. A bulky glycocalyx fosters metastasis formation by promoting G1 cell cycle progression. eLife 6, e25752, doi: 10.7554/eLife.25752 (2017).

17 Tinder, T. L. et al. MUC1 Enhances Tumor Progression and Contributes Toward Immunosuppression in a Mouse Model of Spontaneous Pancreatic Adenocarcinoma. The Journal of Immunology 181, 3116–3125, doi: 10.4049/jimmunol.181.5.3116 (2008).

18 Bast, R. C., Jr., Hennessy, B. & Mills, G. B. The biology of ovarian cancer: new opportunities for translation. Nature reviews. Cancer 9, 415–428, doi: 10.1038/nrc2644 (2009).

19 Rahn, J. J., Dabbagh, L., Pasdar, M. & Hugh, J. C. The importance of MUC1 cellular localization in patients with breast carcinoma. Cancer 91, 1973–1982, doi:doi: 10.1002/1097-0142(20010601)91:11<1973::AID-CNCR1222>3.0.C0;2-A (2001).

20 Wang, X., Lan, H., Li, J., Su, Y. & Xu, L. Muc1 promotes migration and lung metastasis of melanoma cells. American journal of cancer research 5, 2590–2604 (2015).

21 Thompson, C. B. et al. Enzymatic depletion of tumor hyaluronan induces antitumor responses in preclinical animal models. Molecular cancer therapeutics 9, 3052–3064, doi: 10.1158/1535-7163.Mct-10-0470 (2010).

22 Hingorani, S. R. et al. Phase Ib Study of PEGylated Recombinant Human Hyaluronidase and Gemcitabine in Patients with Advanced Pancreatic Cancer. Clinical cancer research: an official journal of the American Association for Cancer Research 22, 2848–2854, doi: 10.1158/1078-0432.Ccr-15-2010 (2016).

23 Xiao, H., Woods, E. C., Vukojicic, P. & Bertozzi, C. R. Precision glycocalyx editing as a strategy for cancer immunotherapy. Proceedings of the National Academy of Sciences 113, 10304–10309, doi: 10.1073/pnas.1608069113 (2016).

24 Kong, F., García, A. J., Mould, A. P., Humphries, M. J. & Zhu, C. Demonstration of catch bonds between an integrin and its ligand. The Journal of Cell Biology 185, 1275, doi: 10.1083/jcb.200810002 (2009).

25 Nordenfelt, P., Elliott, H. L. & Springer, T. A. Coordinated integrin activation by actin-dependent force during T-cell migration. Nature Communications 7, 13119, doi: 10.1038/ncomms13119 (2016).

26 Wang, X. & Ha, T. Defining Single Molecular Forces Required to Activate Integrin and Notch Signaling. Science 340, 991, doi: 10.1126/science.1231041 (2013).

27 Li, J. & Springer, T. A. Integrin extension enables ultrasensitive regulation by cytoskeletal force. Proceedings of the National Academy of Sciences 114, 4685, doi: 10.1073/pnas.1704171114 (2017).

28 Nordenfelt, P. et al. Direction of actin flow dictates integrin LFA-1 orientation during leukocyte migration. Nat. Commun. 8, 2047, doi: 10.1038/s41467-017-01848-y (2017).

29 Brockman, J. M. et al. Mapping the 3D orientation of piconewton integrin traction forces. Nat. Methods 15, 115, doi: 10.1038/nmeth.4536 (2017).

30 Zhang, Y. et al. Platelet integrins exhibit anisotropic mechanosensing and harness piconewton forces to mediate platelet aggregation. Proc. Natl. Acad. Sci. U. S. A. 115, 325, doi: 10.1073/pnas.1710828115 (2018).

31 Kechagia, J. Z., Ivaska, J. & Roca-Cusachs, P. Integrins as biomechanical sensors of the microenvironment. Nature Reviews Molecular Cell Biology 20, 457–473, doi: 10.1038/s41580-019-0134-2 (2019).

32 Sun, Z., Costell, M. & Fässler, R. Integrin activation by talin, kindlin and mechanical forces. Nature Cell Biology 21, 25–31, doi: 10.1038/s41556-018-0234-9 (2019).

33 Bell, G. I. Models for the specific adhesion of cells to cells. Science 200, 618–627 (1978).

34 Atilgan, E. & Ovryn, B. Nucleation and Growth of Integrin Adhesions. Biophys. J. 96, 3555–3572, doi: 10.1016/j.bpj.2009.02.023 (2009).

35 Sarvestani, A. S. The effect of substrate rigidity on the assembly of specific bonds at biological interfaces. Soft Matter 9, 5927–5932, doi: 10.1039/C3SM00036B (2013).

36 Samadi-Dooki, A., Shodja, H. M. & Malekmotiei, L. The effect of the physical properties of the substrate on the kinetics of cell adhesion and crawling studied by an axisymmetric diffusion-energy balance coupled model. Soft Matter 11, 3693–3705, doi: 10.1039/C5SM00394F (2015).

37 Asaro, R. J., Lin, K. & Zhu, Q. Mechanosensitivity Occurs along the Adhesome’s Force Train and Affects Traction Stress. Biophys. J. 117, 1599–1614, doi: 10.1016/j.bpj.2019.08.039 (2019).

38 Oria, R. et al. Force loading explains spatial sensing of ligands by cells. Nature 552, 219, doi: 10.1038/nature24662 (2017).

39 Elosegui-Artola, A. et al. Rigidity sensing and adaptation through regulation of integrin types. Nature Materials 13, 631–637, doi: 10.1038/nmat3960 (2014).

40 Bidone, T. C., Skeeters, A. V., Oakes, P. W. & Voth, G. A. Multiscale model of integrin adhesion assembly. PLoS Comput. Biol. 15, e1007077, doi: 10.1371/journal.pcbi.1007077 (2019).

41 MacKay, L. & Khadra, A. The bioenergetics of integrin-based adhesion, from single molecule dynamics to stability of macromolecular complexes. Computational and Structural Biotechnology Journal 18, 393–416, doi: https://doi.org/10.1016/j.csbj.2020.02.003 (2020).

42 blanchard, a. Burnt bridge ratchet motor force scales linearly with polyvalency: a computational study. Soft Matter, doi: 10.1039/D1SM00676B (2021).

43 Blanchard, A. T. & Salaita, K. Multivalent molecular tension probes as anisotropic mechanosensors: concept and simulation. Phys. Biol. 18, 034001, doi: 10.1088/1478-3975/abd333 (2021).

44 Thomas, W. et al. Catch-Bond Model Derived from Allostery Explains Force-Activated Bacterial Adhesion. Biophys. J. 90, 753–764, doi: https://doi.org/10.1529/biophysj.105.066548 (2006).

45 Helfrich, W. Elastic Properties of Lipid Bilayers: Theory and Possible Experiments. Zeitschrift für Naturforschung C 28, 693–703, doi:doi: 10.1515/znc-1973-11-1209 (1973).

